# Benchmarking Next-Generation Sequencing Platforms: A Comprehensive Comparison Of Single-Cell RNA-Seq from Ultima UG 100 vs. Illumina NovaSeq X Plus

**DOI:** 10.64898/2026.01.16.699571

**Authors:** Ruoyun Xiong, Jaymala Patel, Tamas Sztanka-Toth, Enes Senel, Karl Calara-Nielsen, Yashoda Rajpurohit, Tom Altenburg, Joel Greshock, Denis Smirnov, Alex Javidi

## Abstract

Single-cell RNA sequencing (scRNA-seq) has become an essential technology for dissecting cellular heterogeneity in complex biological systems, with Illumina platforms serving as the dominant NGS provider^1–3^. The recent emergence of the Ultima Genomics UG 100 platform offers a high-throughput and potentially more cost-effective alternative, warranting a rigorous comparative analysis. Here, we conducted a comprehensive head-to-head comparison of the Ultima Genomics UG 100 and Illumina NovaSeq X Plus platforms, generating deep-coverage scRNA-seq data from human PBMCs and stimulated T-cell samples. Our multi-level analysis revealed highly concordant performance in both cell type identification and gene expression. We found that both platforms robustly captured all major immune lineages and demonstrated good transcriptional agreement. Minor observed discrepancies were primarily correlated with low UMI counts, suggesting stochastic biological effects rather than technical variance. Overall, our study establishes that the Ultima UG 100 delivers performance highly comparable to the Illumina NovaSeq X Plus for single-cell transcriptomics.

## Introduction

Single-cell RNA sequencing (scRNA-seq) has revolutionized biological and clinical research, offering unprecedented resolution to dissect cellular heterogeneity and map the complex ecosystems underlying health and disease^1–3^. For over a decade, Illumina’s Sequencing by Synthesis (SBS) technology, with its paired-end, fixed-length read architecture, has been the indispensable gold standard, forming the backbone of global genomics research. However, the cost of sequencing is always a consideration, particularly for large-scale studies where it can become a significant barrier; consequently, more affordable options have recently been emerging in this area^4,5^.

To address this need for scalable and cost-effective solutions, Ultima Genomics has introduced a platform leveraging a distinct, flow-based “mostly natural sequencing-by-synthesis” (mnSBS) chemistry, where nucleotides are introduced sequentially rather than simultaneously, and a single-end, variable-length read architecture^4^. The fundamental differences between Ultima and Illumina technologies necessitate rigorous, unbiased validation to ensure that data from this emerging platform is consistent with the vast body of existing Illumina-generated data. Without such validation, the integration of new technologies into established research pipelines risks introducing confounding technical artifacts that could jeopardize the integrity of longitudinal studies and cross-cohort meta-analyses.

While initial studies have shown promising comparability^6–8^, several critical gaps remain. First, both platforms have continued to evolve, and a direct comparison of the latest-generation instruments is needed. Second, the performance of these platforms in resolving highly similar and challenging cell subtypes, such as those within the T-cell lineage, has not been deeply investigated. Finally, cross-validating platform performance with diverse biological samples is essential to build a robust evidence base for the broader research community.

Here, we perform a direct, head-to-head comparison of the latest high-throughput instruments, the Ultima UG 100 and the Illumina NovaSeq X Plus. We designed a study to systematically evaluate every stage of a typical scRNA-seq workflow, from foundational quality control metrics to high-level biological interpretation. To achieve this, we prepared five 10x Genomics single-cell libraries from clinically relevant stimulated T cells (n=3) and PBMCs (n=2), splitting each library for parallel sequencing on both platforms to minimize technical variability (Figure 1a). By applying a comprehensive analytical framework, we interrogated concordance at the level of individual cells, cell populations, and individual genes. Our findings provide a critical validation of the Ultima platform, offering the evidence needed to guide its adoption for future large-scale single-cell genomics initiatives.

**Figure 1:**
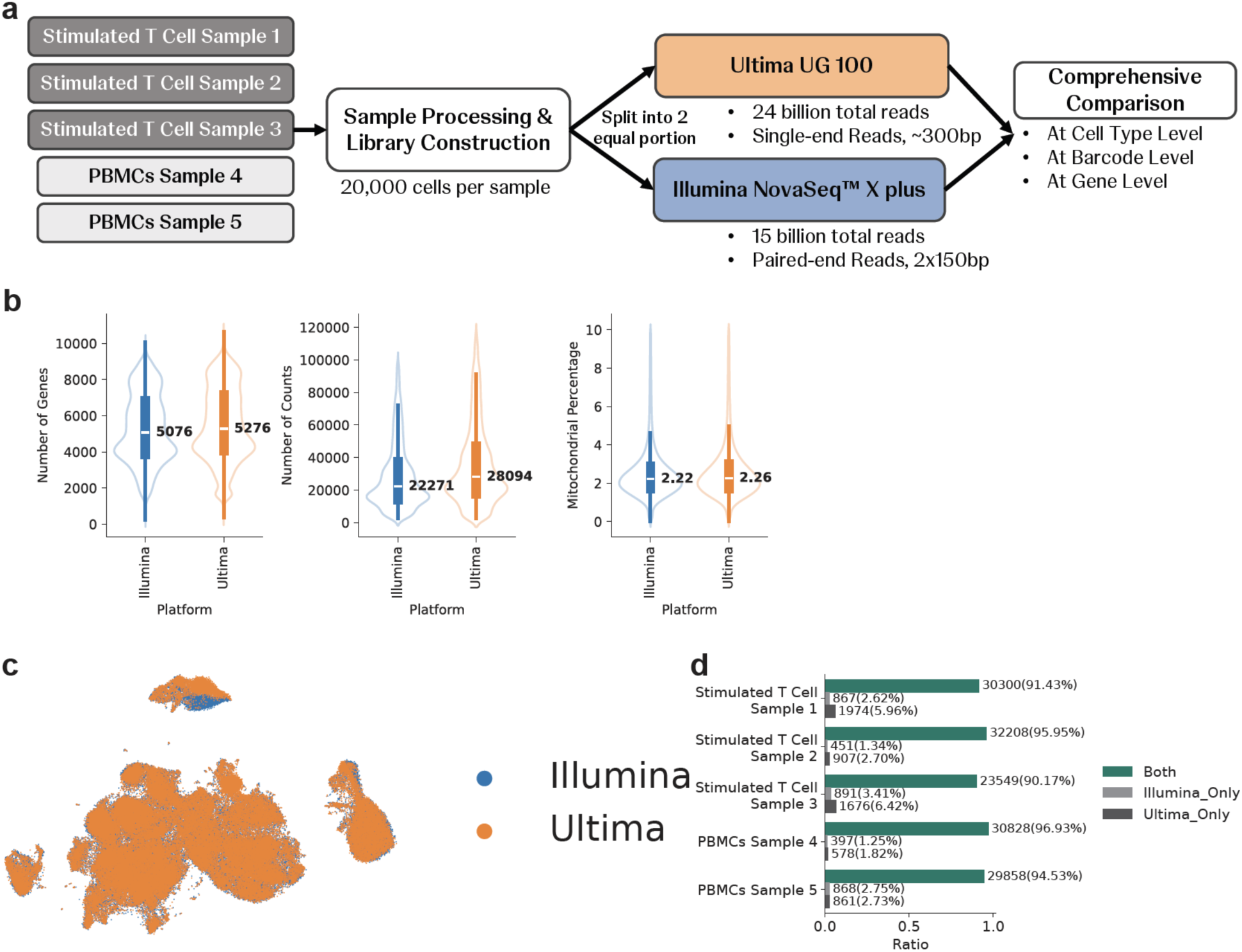
Study Workflow and Comparison of Sequencing Quality Metrics. This figure outlines the experimental design for comparing the Ultima and Illumina platforms and presents key quality control (QC) metrics and barcode detection overlap. **a) Experimental Design Workflow.** Five 10x Genomics libraries were prepared: three from stimulated T-cell samples and two from peripheral blood mononuclear cell (PBMC) samples, each containing approximately 20,000 cells. Each library was split into two equal portions for parallel sequencing. One portion was sequenced on the Illumina NovaSeq™ X plus, generating 15 billion total paired-end reads (2×150bp). The other portion was sequenced on the Ultima UG 100, generating 24 billion total single-end reads (∼300bp). **b) Comparison of Key Quality Control Metrics.** Violin plots compare the per-cell distributions of three QC metrics between the Illumina (blue) and Ultima (orange) platforms. The width of each violin illustrates the density of cells at a given value, with the internal box plot showing the median and interquartile range. **c) Barcode Detection Overlap Between Platforms.** Stacked bar charts quantify the concordance of cell barcode identification for each of the five samples. Each bar is segmented to show the proportion of barcodes detected exclusively on the Illumina platform (light grey), exclusively on the Ultima platform (dark grey), or on both platforms (green). **Abbreviations:** PBMC, Peripheral Blood Mononuclear Cell; UMI, Unique Molecular Identifier.

## Results

### Comparable Overall Performance and Quality Metrics Across Platforms

We first assessed key quality control (QC) metrics from five paired samples to establish a baseline for platform comparison. Our analysis revealed that both the Ultima UG 100 and the Illumina NovaSeq™ X Plus demonstrated highly comparable performance. We examined three primary indicators of library quality: the number of genes detected per cell^9^, the number of Unique Molecular Identifiers (UMIs) per cell^10^, and the percentage of mitochondrial reads^11^. The median number of genes detected per cell was similar between platforms, with 5,076 for Illumina and 5,276 for Ultima. We observed a 26% increase in median UMI counts for Ultima (28,094) compared to Illumina (22,271), which is consistent with the greater sequencing depth allocated to the Ultima runs. Critically, the percentage of mitochondrial reads, a key indicator of cell stress^11,12,13^, was nearly identical between Illumina (2.22%) and Ultima (2.26%), demonstrating that neither platform introduces a bias toward lower-quality cells (Figure 1b).

To evaluate the global similarity between the datasets, we performed Uniform Manifold Approximation and Projection (UMAP) dimensionality reduction^14,15^. The resulting plot showed a high degree of integration, with cells from both Illumina and Ultima intermixing and for most clusters, there are no evidence of platform-driven distinct (Figure 1c), except one cluster on the top, which we found the key driver in the later content (Figure 4e). This indicates that the biological variance captured in the transcriptomes outweighs the technical variance introduced by the sequencing platforms (Supplemental Table 1).

Furthermore, we confirmed that most cell barcodes were successfully identified in both sequencing runs for each of the five physically split libraries. On average, over 90% of all cellular barcodes were shared between the paired Illumina and Ultima datasets for each sample (Figure 1d). To ensure the most direct and fair comparison, all subsequent analyses were performed exclusively on this set of shared barcodes, guaranteeing that our downstream comparisons are based on the exact same populations of single cells.

### Ultima and Illumina Showed High Concordance in Immune Cell Composition

To determine if Ultima and Illumina sequencing resolve the same cellular subpopulations at similar proportions, we performed an integrated clustering analysis (using scVI) on the combined dataset of shared barcodes (Methods). After assigning cell type identities using automated annotation models by canonical immune markers (CellTypist, see Methods), our analysis revealed a high degree of concordance between the two platforms. At a broad level, we identified the major expected immune lineages, including T cells, B cells, monocytes, dendritic cells (DCs), and innate lymphoid cells (ILCs) (Supplemental Figure 1a). T cells were further resolved into stimulated and non-stimulated populations. When we examined the contribution of each platform to these cell type clusters, we found a consistent and unbiased split; for every major cell type, the population was composed of approximately 50% cells from Illumina and 50% from Ultima (Figure 2a-b). This balanced representation indicates no platform-specific bias in the capture of broad cellular categories, a finding that holds for both common populations like T cells and rarer populations like DCs.

**Figure 2:**
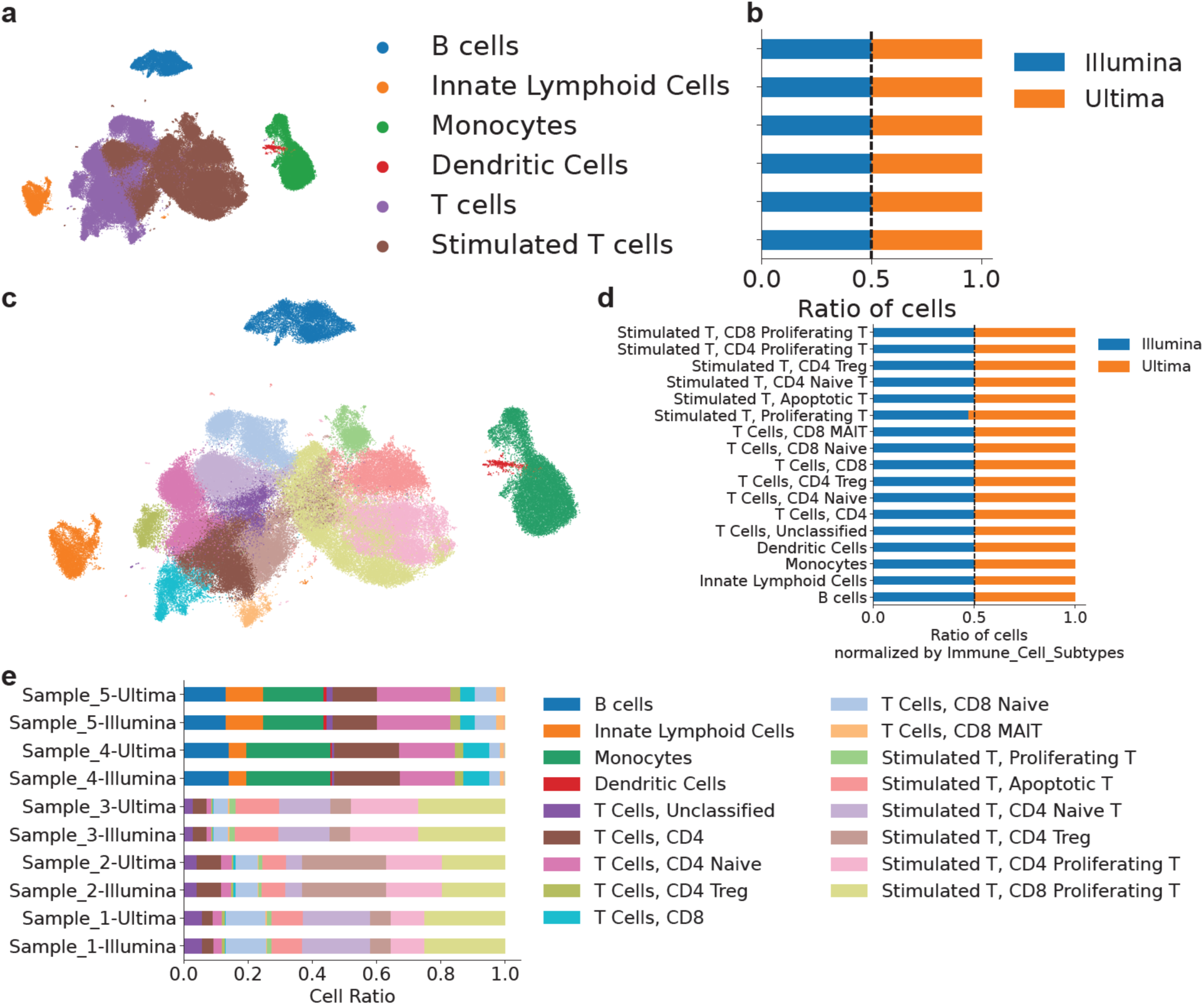
High Concordance of Immune Cell Type Composition Between Platforms. This figure demonstrates the consistency of cell type identification and proportional representation between the Illumina and Ultima sequencing platforms at both broad and granular resolutions. **a-b) Proportional Contribution to Major Immune Lineages.** UMAP and stacked bar charts show the six major immune cell populations identified across five samples and the relative contribution of cells sequenced on the Illumina (blue) and Ultima (orange) platforms. The x-axis represents the ratio of cells, with a dashed vertical line at 0.5 indicating an equal 50/50 split between the two platforms. Each horizontal bar corresponds to a major cell type, including B cells, monocytes, and T cells. **c-d) Proportional Contribution to Granular Immune Subtypes.** This panel extends the analysis to 17 distinct immune cell subtypes resolved from a more granular sub-clustering with a specific focus on T cell populations. The stacked bar charts again show the proportional contribution of Illumina (blue) and Ultima (orange) to each subtype. **e) Conservation of Cell Subtype Proportions Within Biological Samples.** Stacked bar plots compare the detailed cell type composition for each of the five biological samples. For each sample, the proportional representation from the Ultima run (top bar) is paired with the corresponding Illumina run (bottom bar). The x-axis represents the cell ratio, with the total composition of each run summing to 1.0. Each colored segment corresponds to one of the 17 immune subtypes identified. **Abbreviations:** UMAP, Uniform Manifold Approximation and Projection.

We then performed a more granular sub-clustering analysis, which resolved 17 distinct immune cell subtypes, including 13 T cell subtypes that spanned naive, effector, memory, and regulatory populations^16–21^ (Supplemental Figure 1a-b). Even at this higher resolution, the platform concordance remained high. Most subtypes, such as, CD8 Naive T Cells and Stimulated CD4 Treg Cells, maintained the 50/50 split between Ultima and Illumina (Figure 2c-d, Supplemental Table 1). Furthermore, the proportional representation of these 17 subtypes was highly consistent between the paired Illumina and Ultima runs for each of the five biological samples (Figure 2e, Supplemental Table 1). This robust conservation of cellular composition, from broad lineages down to fine-grained subtypes, demonstrates that both technologies provide a similar resolution for the complex cellular architecture of biological tissues. Our results suggest that biological conclusions about cell population dynamics from one platform would be recapitulated on the other.

### Low Read Counts Drive Discrepancies in Cell Type Similarity

To assess transcriptomic similarity at the single-cell level, we quantified the agreement between platforms for all shared barcodes by calculating the pairwise Pearson correlation coefficient between their Illumina– and Ultima-derived gene counts. This analysis revealed a high overall correlation across all cells, signifying a strong agreement in gene expression quantification at the single-cell level (Figure 3a, Supplemental Table 1).

**Figure 3:**
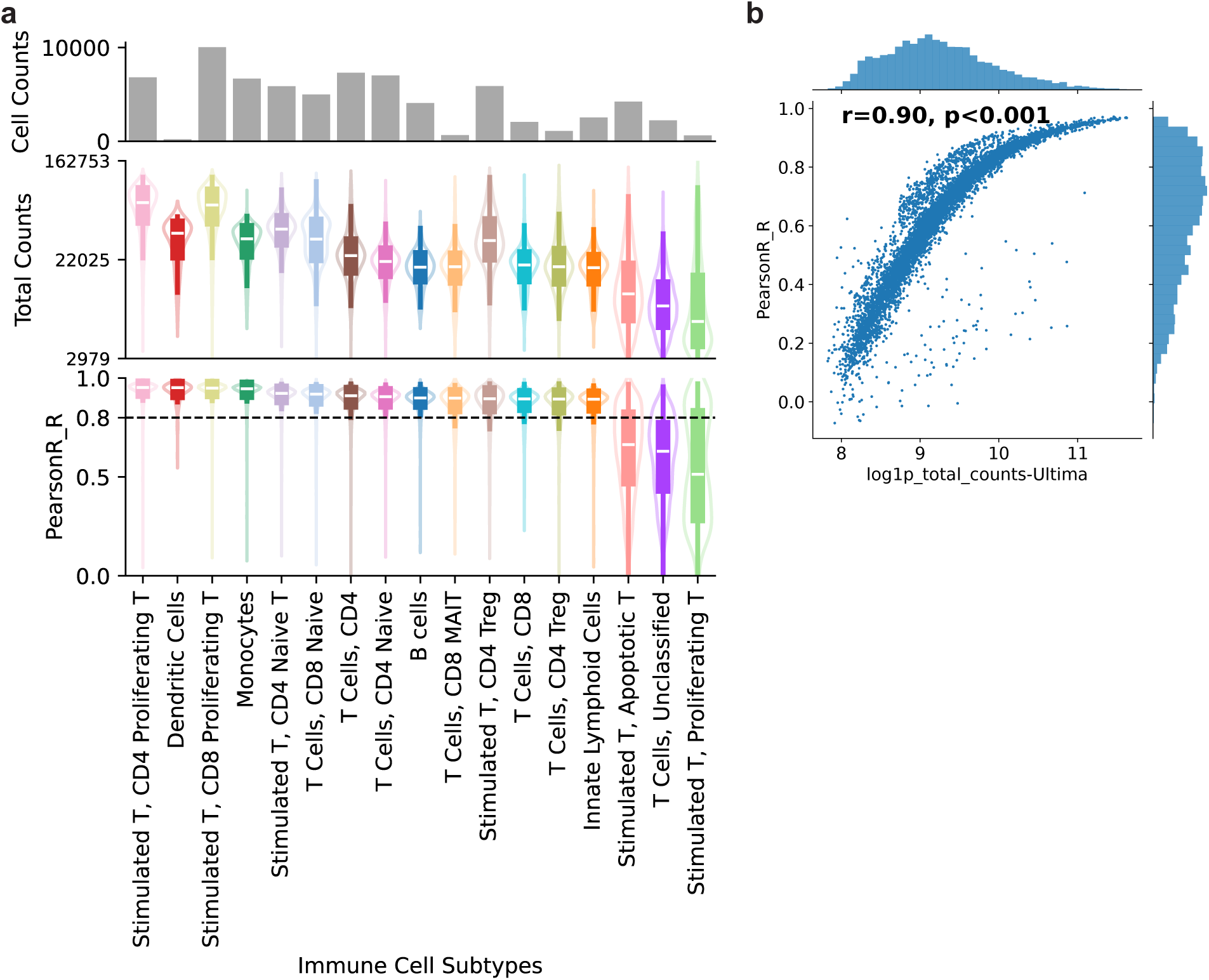
Single Cell Level Similarity Between Platforms is Correlated with Read Counts. This figure illustrates the relationship between per-cell transcriptomic similarity (Pearson R) and total UMI counts across 17 identified immune cell subtypes. **a) Per-Cell Transcriptomic Correlation Across Immune Subtypes.** This multi-panel plot displays three metrics for each of the 17 immune cell subtypes, which are ordered along the shared x-axis by their median Pearson correlation coefficient. **Top panel**, The total number of cells within each subtype is shown as a bar chart (y-axis: Cell Counts). **Middle panel**, The distribution of total UMI counts per cell is illustrated with violin plots (y-axis: Total Counts). The width of each violin represents the density of cells at a given UMI count, with the internal box plot showing the median and interquartile range. **Bottom panel**, The distribution of per-cell Pearson correlation coefficients (R) between paired Illumina and Ultima transcriptomes is shown with violin plots (y-axis: Pearson R). The dashed horizontal line is set at R=0.8 for reference. **b) Correlation Between Read Counts and Transcriptomic Similarity.** This scatter plot shows the direct relationship between cellular read counts and platform concordance for cells from the most dissimilar subtypes (Stimulated T, Apoptotic T; T Cells, Unclassified; and Stimulated T, Proliferating T). Each blue dot represents a single cell. The x-axis represents the log-transformed total UMI counts from Ultima, and the y-axis represents the per-cell Pearson R value comparing the Ultima and Illumina data for that cell. **Abbreviations:** R, Pearson Correlation Coefficient; UMI, Unique Molecular Identifier.

We then dissected this correlation by examining its distribution within each of the 17 identified immune cell subtypes. We found that the majority of cell types exhibited a high degree of transcriptomic similarity, with median Pearson R values consistently above 0.8 (Figure 3a, bottom panel). However, we observed that three T-cell subtypes, namely Stimulated Apoptotic T Cells, Unclassified T Cells and Stimulated Proliferating T Cells, displayed significantly lower and more variable correlation scores. A closer examination revealed that these specific subtypes were also characterized by having the lowest median UMI counts per cell on both Illumina and Ultima platforms. (Figure 3a, middle panel).

This relationship suggests that the observed discrepancies are not due to systematic technological bias but rather reflect the fundamental challenge of stochastic gene capture in cells with low RNA content. We statistically confirmed this link by correlating the per-cell UMI counts with the per-cell Pearson R values for these three dissimilar subtypes. This analysis showed a strong, positive correlation (R=0.90, p<0.001), demonstrating that higher cellular RNA content directly predicts higher transcriptomic concordance between the two platforms (Figure 3b).

### Comparable Gene Expression Profiles Across Ultima and Illumina Platforms

Finally, we systematically compared the platforms’ performance at the level of individual gene expression. To test whether differences were driven by technical or biological factors, we first created pseudo-bulk expression profiles by aggregating UMI counts for each gene across all cells^22–24,25^. This analysis revealed a significant concordance, with 99.08% of approximately 13,000 highly expressed genes (defined as having minimum counts >10 and total counts >50 per sample) showing no significant difference in expression between the two platforms (Figure 4a-c). Only 120 genes, representing less than 1% of the expressed transcriptome, were identified as differentially expressed, with 72 genes showing higher expression in Illumina data and 48 in Ultima data (Figure 4b).

**Figure 4:**
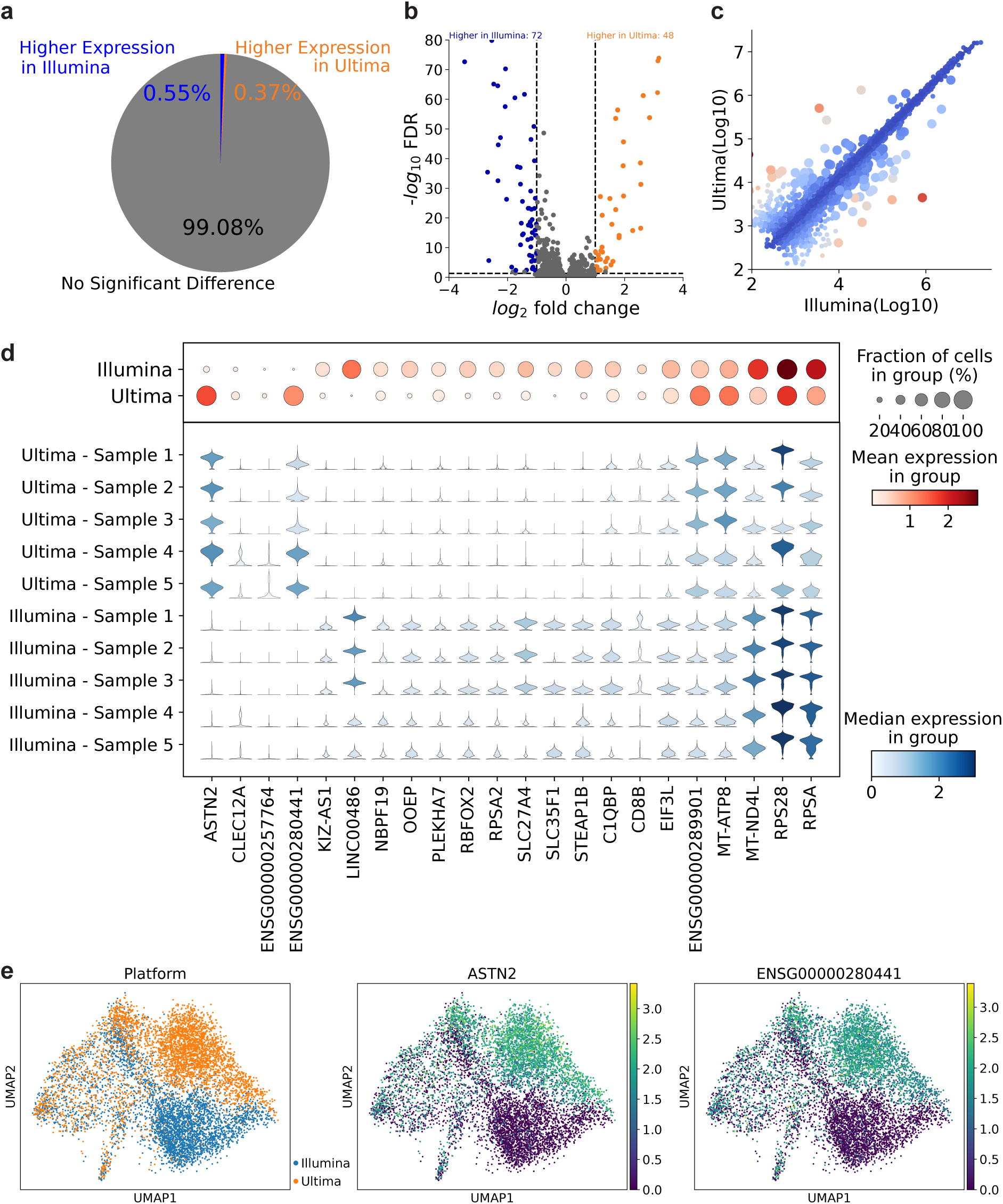
Gene Expression Level Comparison of Ultima and Illumina. This figure compares pseudo-bulk and single-cell gene expression data to demonstrate the high level of agreement between the Illumina and Ultima sequencing platforms. **a) Summary of Differentially Expressed Genes.** This pie chart visually summarizes the differential expression analysis from panel A. It highlights that the proportion of genes with higher expression in Illumina (0.55%) and Ultima (0.37%) together account for less than 1% of the total genes, with the remainder (99.08%) showing no significant difference. **b) Correlation of Gene Expression.** This scatter plot compares the log10-transformed the expression level of each gene from the Illumina platform (x-axis) against the Ultima platform (y-axis). Each point represents an individual gene. **c) Volcano Plot of Pseudo-bulk Differential Gene Expression.** This plot compares the pseudo-bulk expression of approximately 13,000 genes between the two platforms. The x-axis represents the log_2_fold change, and the y-axis represents the statistical significance as the –log_10_ FDR, representing the adjusted p-value. Genes with no significant difference in expression are colored gray. A small number of genes were identified as significantly more highly expressed in the Illumina data (blue, n=72) or the Ultima data (red, n=48). **d) Expression of Differentially Expressed Genes Across Samples.** This dot plot displays the expression patterns for a selection of the most significantly differentially expressed genes (x-axis) across each of the 10 individual sequencing runs (y-axis). The size of each dot indicates the fraction of cells within that sample expressing the gene, while the color intensity represents the mean expression level within the expressing cells. **e) UMAP Visualization of a Platform-Specific Batch Effect.** These three UMAP plots visualize gene expression specifically within the B-cell population. **Left panel**, Cells are colored by sequencing platform (Illumina, orange; Ultima, blue) **Middle and Right panels**, Cells are colored by the normalized expression level of two differentially expressed genes, *ASTN2* and *ENSG00000280441*. The plots show that these genes are almost exclusively expressed in the cells sequenced on the Ultima platform (purple), identifying these genes as the primary drivers of the minor batch effect observed in the left panel. **Abbreviations:** UMAP, Uniform Manifold Approximation and Projection; FDR, False Discovery Rate.

To investigate the origin of these differences, we examined their consistency across samples. We found that the expression patterns of key differentially expressed genes, including several ribosomal and mitochondrial genes, are highly consistent across all five samples from both PBMC and Stimulated T Cell samples (Figure 4d, Supplemental Figure 2). This lack of inter-tissue variability suggests that these minor differences are attributable to technical characteristics of the sequencing platforms rather than biological variation^26,27^.

We next assessed the impact of these technical variations on downstream biological interpretation. Previously, we found a batch effect localized within the B-cell cluster (Figure 1c, 4e). A targeted analysis revealed that this platform-driven separation was attributable to the specific detection of two genes, *ASTN2*^28–30^ and the uncharacterized locus *ENSG00000280441*, by the Ultima platform (Figure 4e). These genes were robustly detected in B cells sequenced on Ultima but were virtually absent in the same cells sequenced on Illumina across all samples (Figure 4d). While the precise technical reason for this discrepancy is unclear, it may relate to differences in how the longer, single-end Ultima reads map to these specific transcripts compared to paired-end Illumina reads^31,32, 33^.

## Discussion

The increasing scale of biomedical research necessitates sequencing technologies that are both high-quality and cost-effective. Our study provides a direct comparison of the latest high-throughput single-cell sequencing platforms from Ultima and Illumina, establishing a critical benchmark for the research community^,35^. We demonstrate that data generated from both platforms are highly comparable across key metrics, from the identification of broad and fine-grained cell populations to the quantification of the vast majority of individual genes^36–38^. These findings provide an essential validation, indicating that large-scale single-cell studies can be performed with both platforms without compromising core biological conclusions^39^.

A primary goal of this study was to understand the source of any observed differences between the platforms. Our results demonstrate high concordance between Ultima and Illumina for scRNA-seq, aligning with the findings of other platform comparisons^6,7^. More importantly, our work extends these findings in several key aspects. First, we report robust performance in resolving 13 transcriptionally similar T-cell subtypes—a challenging use case not deeply explored in prior work but critical to many immunology and oncology applications^16,40,41^. Second, we found that lower transcriptomic correlation between individual cells was directly linked to low UMI counts. This is not a platform-specific bias but rather reflects the inherent stochasticity of gene capture in cells with low RNA content, a fundamental challenge in scRNA-seq and NGS applications. Finally, we identified rare, technology-specific artifacts. The minor batch effect within the B-cell cluster, for example, was attributable to the differential detection of just two genes, *ASTN2* and *ENSG00000280441*.^42–44^ While the precise mechanism is unknown, we hypothesize that this is due to the fundamental differences in read architecture between the platforms. The longer, single-end reads from Ultima may map more efficiently to these specific transcripts than Illumina’s paired-end reads, particularly if the distinguishing sequences lie at the transcript ends^6,45,46^.

This analysis provides a clear, data-driven framework for researchers when choosing a sequencing platform. The decision should be guided by the specific goals of the experiment^47,48^. For large-scale, discovery-oriented projects where maximizing sample number is critical for statistical power, the cost efficiency of the Ultima platform is a significant advantage^33^. In contrast, for studies that require maximal consistency with large legacy datasets generated on Illumina platforms, or for those focused on specific biological questions involving genes known to be rare artifacts, the incumbent technology may be more appropriate. Furthermore, applications that depend on paired-end read information, such as alternative splicing analysis, remain a key domain for Illumina’s architecture^49,50^.

Limitations of our study include its focus on blood-derived samples (PBMCs and stimulated T-cell subtypes). While this comparison provides a critical baseline, future work is needed to extend this validation across a wider range of tissues complexities and sample qualities. For example, for solid tumor profiling, where high cellular heterogeneity challenges precise subtype definition, future benchmarks could employ deeper sequencing depths or integrate orthogonal validation methods, such as histology or spatial transcriptomics. Such studies will contribute to a more complete understanding of each platform’s performance envelope and help drive the development of a mature bioinformatic ecosystem, with tools specifically optimized for the unique data characteristics of each technology^51–54^.

## Methods

### Single-Cell Library Preparation

CITE-seq libraries were prepared from stimulated T cells (T cell activation via CD3 and CD28 with T Cell TransAct– # milteyni Biotech 130-111-160^55^) and unstimulated PBMCs using the Chromium GEM-X Single Cell 3’ v4 chemistry^56^ (UserGuide_CG000731RevB). Following the user guide, cDNA was amplified from 20,000 cells. Final gene expression (GEX) libraries were constructed in duplicate, generating two sequencing-ready replicates. One replicate was sequenced on the Ultima UG100 platform (achieving ∼240,000 reads per cell), and the other on the Illumina NovaSeq X Plus platform (∼150,000 reads per cell) at CRO (Referance ELN E259195 for details. Comprehensive analyses were performed on data from both platforms to compare cell type distributions, barcode assignments, and gene-level expression profiles.

### Raw Data Pre-processing and Alignment

Raw base call (BCL) files from both the Ultima and Illumina sequencing platforms were demultiplexed and converted to FASTQ format. The resulting FASTQ files were processed using Cell Ranger (v7.1.0; 10x Genomics^57^). Reads were aligned to the GRCh38 reference genome^58^, and Unique Molecular Identifiers (UMIs) were counted to generate cell-by-gene count matrices for each sample.

### Quality Control and Data Filtering

Subsequent quality control (QC) was performed on the raw count matrices using scanpy (v1.10.4)^59^. For each sample, cells were filtered based on several criteria. First, we removed cells where the log-transformed gene counts or UMI counts were more than 3 median absolute deviations (MADs) from the sample median. Additionally, cells with a mitochondrial gene content exceeding 10% were discarded. Potential doublets were identified using Scrublet^60^, and any cell with a doublet score greater than 0.3 was removed. Finally, we filtered out genes that were expressed in fewer than 20 cells across the entire dataset.

### Data Integration, Cell Clustering and Visualization

To mitigate batch effects arising from different samples, the filtered count matrices were integrated using the Single-Cell Variational Inference (scVI^61^) model from the scvi-tools package (v1.3.2). Cell clustering was performed on the scVI-generated latent embedding. A shared nearest neighbor (SNN) graph was constructed based on the Euclidean distance in the latent space, and the Leiden community detection algorithm was applied to identify distinct cell clusters (scanpy.tl.leiden). The clustering resolution was set to 0.6 to achieve an optimal separation of cell populations. For visualization, Uniform Manifold Approximation and Projection (UMAP) was run on the scVI latent representation to generate a two-dimensional embedding of the data.

### Automated Cell Type Annotation

To assign biological identities to the cell clusters, we employed a hierarchical annotation strategy using celltypist^62^ (v1.7.1). First, broad immune lineages were identified using the Immune_All_High model. These coarse-grained labels were then refined into more granular subtypes using the Immune_All_Low model (https://www.celltypist.org/models)^63^. Final cell type annotations were assigned based on the highest confidence score from the most specific applicable model.

### Shared Barcode Filtering

To ensure a direct 1-to-1 comparison, the filtered barcode-gene matrices from each paired Illumina and Ultima run were filtered to retain only the intersection of their respective cell barcodes. All subsequent analyses were performed exclusively on this set of shared barcodes, guaranteeing that all comparisons are made on the-exact same population of single cells.

### Transcriptomic Correlation Analysis (Barcode-by-Barcode)

To assess the concordance at the single-cell level, we performed a direct “barcode-by-barcode” correlation. This analysis assumes the same 10x Genomics library was split and sequenced on both platforms, resulting in a shared set of cell barcodes.

For each individual matched cell barcode *i*, we extracted its log-transformed (log1p) gene expression vector from the Ultima platform (*U_i_*) and the Illumina platform (*I_i_*). We then computed the Pearson correlation coefficient (*r*) between these two vectors.

The Pearson correlation coefficient *r* for any two vectors *x* (e.g., *U_i_*) and *y* (e.g., *I_i_*) of length *n* (number of genes) is defined as:

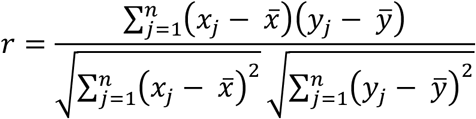

where:

- *n* is the total number of genes.
- *x_j_* and *y_j_* are the individual log-transformed expression values for gene *j* in the Ultima and Illumina vectors, respectively.
- 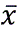 and 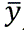 are the mean expression values of vectors *x* and *y*, respectively.

This calculation was performed for all matching barcodes, and the resulting distribution of *r* values was used to assess per-cell transcriptomic concordance between the platforms (Figure 3a).

### Differential Gene Expression (DGE) Analysis

To compare gene expression at a pseudo-bulk level, we first aggregated the raw UMI counts for each gene across all shared barcodes, creating one pseudo-bulk profile for the Ultima platform and one for the Illumina platform. We performed differential expression analysis on these pseudo-bulk count vectors using pydeseq2^63^ (v0.5.0), a Python implementation of the DESeq2 method. The model design was ∼ platform, testing for gene expression changes attributable to the sequencing platform. Genes were considered differentially expressed if the False Discovery Rate (FDR) was less than 0.05 (Figure 4a-c).

## Data and Code

For information about the data and code supporting this study, please contact the corresponding author.

## Acknowledgement and Funding

This work represents a collaborative effort between the Data Science and Oncology Translational Research teams. We thank Brad Foulk and Gourav Choudhary to support Technology introductions review, experimental samples generation and Data Analysis review.

## Author Contributions

Conceptualization: AJ, DS, JG; Coding: RX, TST, ES, TA; Data Curation: TST, ES, AJ; Formal Analysis: RX; Sample design and collection: JP, DS; Investigation: JP, DS, KN, YR, RX, TST, AJ; Visualization: RX, TST, ES; Project Administration: AJ, DS, JG; Supervision: JG, DS, AJ; Resources: JG, DS, AJ; Writing and Editing: RX, JP, TST, ES, AJ, DS

## Lead Contact

Further information and requests for resources and reagents should be directed to the lead contact, Alex Javidi.

## Supplemental Materials

**Supplemental Figure 1:**
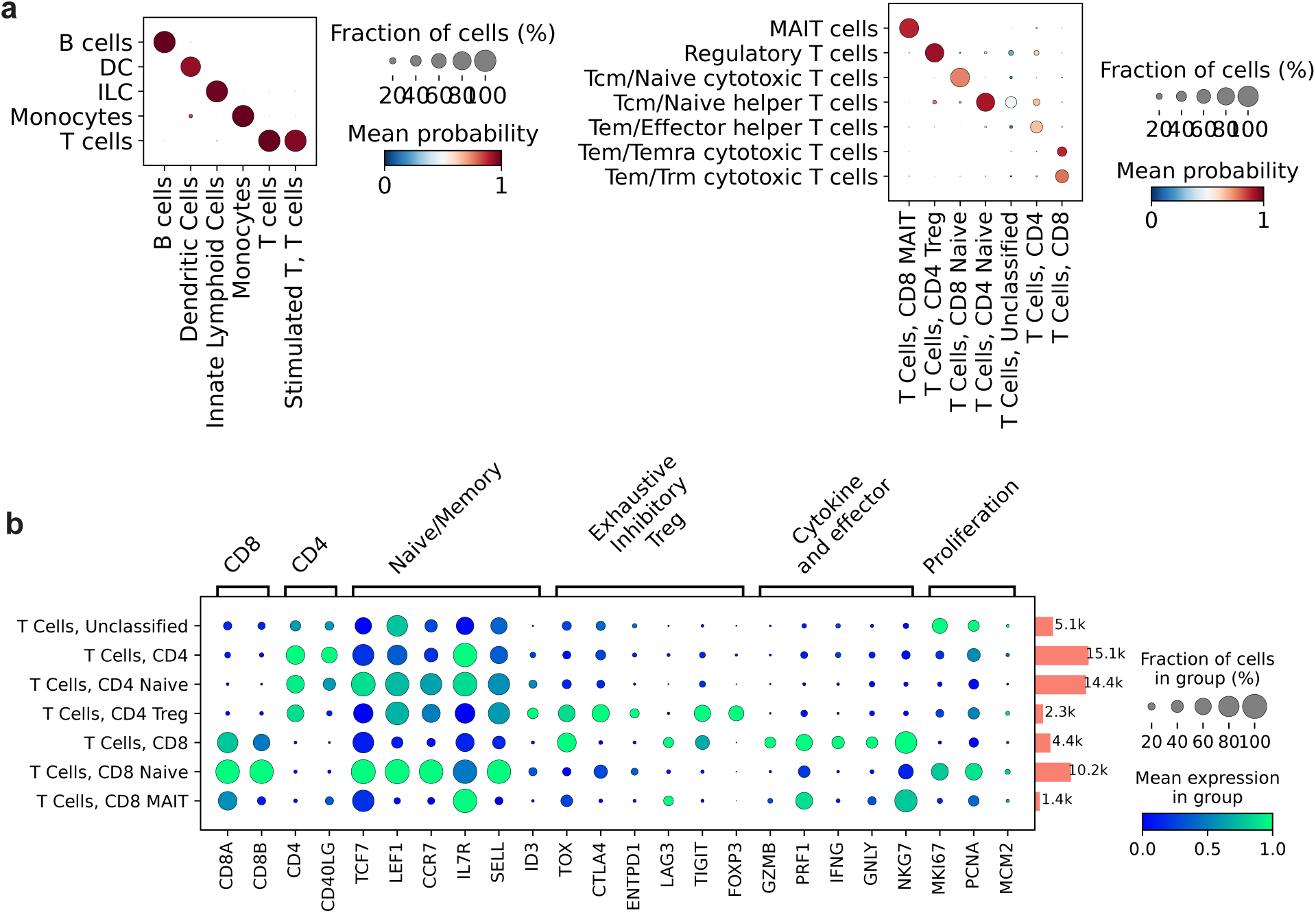
Cell Type Annotation Probability and Marker Gene Validation. This figure provides supporting data for the cell type annotation process, showing the classification probabilities from the automated cell typing tool and validating the T cell subtype annotations with canonical marker gene expression. **a) Cell Type Classification Probabilities.** Dot plots display the probability scores from Cell Typist (Method). For each predicted cell type, the color indicates the mean probability score of the assignment, and the dot size represents the fraction of cells assigned to that type. **b) Canonical Marker Gene Expression for T Cell Subtypes.** This dot plot validates the T cell subtype annotations by displaying the expression of key canonical marker genes grouped by function (e.g., CD4, CD8, Naive/Memory, Proliferation). Each row corresponds to a T cell subtype, and each column represents a gene. The size of each dot is proportional to the percentage of cells within a subtype expressing the gene, while the color intensity corresponds to the average expression level within those cells.

**Supplemental Figure 2:**
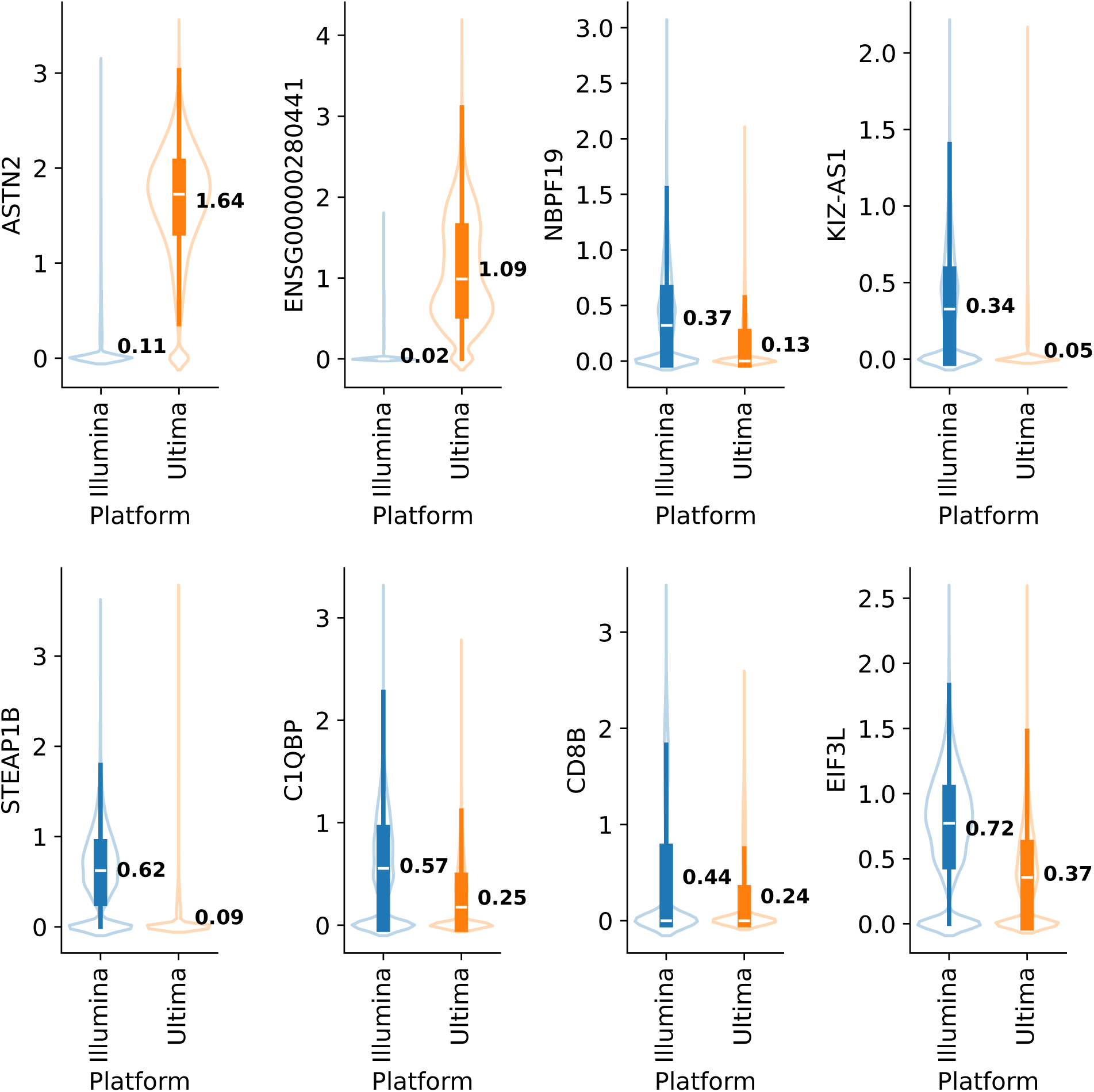
Expression Distributions of Top Differentially Expressed Genes. This figure provides a detailed view of the single-cell expression patterns for eight of the most significant genes identified as differentially expressed between the Illumina and Ultima platforms. For each gene, the distribution of expression values for cells sequenced on the Illumina platform (blue) is compared to those from the Ultima platform (orange). The width of each violin illustrates the density of cells at a given expression level, and the mean expression value is annotated for each platform (e.g., for ASTN2, the mean is 0.11 for Illumina and 1.64 for Ultima).

**Supplemental Table 1:** Cell Counts and Proportions for Immune Subtypes.

